# New models to study plasma cells in mouse based on the restriction of IgJ expression to antibody secreting cells

**DOI:** 10.1101/2020.08.13.249441

**Authors:** Maria Victoria Ayala, Amélie Bonaud, Sébastien Bender, Jean-Marie Lambert, Fabien Lechouane, Claire Carrion, Michel Cogné, Virginie Pascal, Christophe Sirac

## Abstract

Plasma cells (PC) represent the last stage of B cell development and are mainly characterized by their capacity of secreting large quantities of antibodies. They can be implicated in a broad-spectrum of neoplastic disorders, including Multiple Myeloma, Waldenstrom macroglobulinemia or Monoclonal Gammopathy of Clinical Significance, all characterized by the abnormal proliferation of a PC clone. Up to date, there are only few reporter models to specifically follow PC development, migration and homing in mouse and none allowing the genetic manipulation of these cells. We created a transgenic mouse model in which a green fluorescent protein gene was placed under the control of the well-characterized regulatory elements of the murine immunoglobulin J (IgJ) chain locus. Thanks to this model, we demostrated that IgJ is an early and specific marker of antibody secreting cells (ASCs) and appears before the expression of CD138, making it a good candidate to targeted genetic modifications of plasma cells. Therefore, a conditional deletion model using a Tamoxifen-dependent Cre recombinase inserted into the IgJ locus was characterized. Using a reporter model, we showed that, in contrast with existing models of B cell lineage genetic modification, the activity of the CRE recombinase only affects ASCs after tamoxifen treatment. Additionally, we used this model in a functional *in vitro* assay, to show that Ig modifications directly affect plasma cell survival. These two new mouse models, IgJ^GFP^ and IgJ^CreERT2^ represent exquisite tools to study PCs. In pathology, the IgJ^CreERT2^model opens new frontiers for *in vivo* genetic modifications of PCs to better reflect the pathophysiology of PC-related diseases.

## INTRODUCTION

Plasma cells (PC), dedicated to the secretion of large amounts of antigen-specific immunoglobulins, represent the final stage of B cell lineage differentiation. The transcriptional program involved in the transition from B cell to PC has been extensively investigated in the past few years revealing two sets of transcription factors that are mutually opposite: those involved in the B cell identity and maintenance like Pax5, Bcl6 or Bach2 and those that control PC differentiation like Blimp1, Irf4 or Xbp1 ^1–9^. However, few studies make it possible to understand the mechanisms involved after the differentiation, leading to migration, death or on the contrary to the exceptionally long survival of these cells, as well as their deregulation in the case of clonal plasma cell proliferations such as Monoclonal Gammopathy of Undertemined Significance (MGUS) or Multiple Myeloma (MM). One reason is the lack of models to specifically follow PCs in mouse and to turn on/off genes specifically in this lineage. The current murine models of inducible gene invalidation in the B cell lineage only allow deletion at relatively early stages of development (CD19^Cre^, CD21^Cre^). For instance, many models of overexpression of oncogenes normally involved in the development of PC proliferation resulted in general B hyperplasia or B cell tumor developments mainly because the oncogene was expressed throughout B cell development rather than restricted in the plasma cell compartment ^10–12^. Currently, models with germinal center B cell-specific Cre expression (AID^CreERT2^ or IgG1^Cre^) represent the latest stage of B cell differentiation ^13,14^. They can be useful to study the early stages of PC differentiation or transformation but remain useless for fully differentiated PCs. One reason for this lack of PC-specific Cre-deleter model is the difficulty to find a true PC-specific promoter to induce expression of Cre only in PCs. Genes overexpressed in PCs are often expressed earlier during B cell development or in other immune cell lineage (Prdm1, Irf4)^15,16^. Prdm1 promoter-driven reporter genes has been extensively used to decipher PC differentiation and homing^17,18^ since Blimp1 is highly expressed in PCs^19^. However, it was shown that GFP is also expressed in other immune cells^20,21^ precluding the usage of its promoter for a PC-specific Cre-deleter strain. In the absence of such strain, Tellier *et al*. advantageously used the ubiquitous tamoxifen-inducible Rosa26-Cre^ERT2^ strain to study the role of Blimp1 and Ifr4 in mature PCs^22^. However, such approach also induced Cre-driven deletion in other cells expressing these genes and cannot be used to delete or activate genes or oncogenes specifically in PCs.

Immunoglobulin Joining (IgJ)-chain is a 15 kDa peptide expressed by antibody secreting cells (ASCs) and required for correct assembly of pentameric IgM and dimeric IgA ^23^. Several studies have demonstrated that PCs producing monomeric Ig isotypes also express this peptide but it still remains controversial to what extent ^24–26^. The exact stage of production during B cell development and differentiation into PC remains unknown but promoter activity and transcriptional studies have revealed that IgJ is expressed at very early steps of PC differentiation ^27^, before the requirement for Blimp1 over-expression ^28^. Older studies also showed IgJ expression in pre-B and mature B cells^24^.

In the present study, we first created a transgenic murine model that expresses the eGFP under the control of the murine IgJ gene enhancer/promoter ^27^ to follow IgJ expression during *in vivo* and *in vitro* differentiation of B-cells into PC. We show that, in contrast to previous assumptions, almost all ASC express the J-chain, independently of the secreted IgH isotype and its expression is mainly restricted to ASCs with a low expression in GC B cells. In contrast with other PC reporter models based on Prdm1 expression pattern^17,18^, no other hematopoietic lineage yielded GFP expression making this model a unique tool to specifically access PCs localization. IgJ appears as an early marker of terminal differentiation before the expression of CD138, making of it an ideal promoter for a conditional deletion model in PCs. Consequently, we characterized a new model in which a tamoxifen-dependent Cre recombinase (Cre^ERT2^ recombinase) was placed under the control of this promoter in a knock-in transgenic mouse model. For the characterization of its targeted activity, mice were further crossed with a reporter mouse line. This model relies on the expression of a fluorescent protein, tdTomato, after elimination of a floxed STOP cassette. Thanks to it, we were able to show that Cre^ERT2^ recombinase activity is strictly restricted to PCs *in vivo* and *in vitro*. Furthermore, we confirmed this specificity in a model of Heavy Chain Deposition Disease (HCDD) ^29^, yielding a floxed CH1 domain in a transgenic human heavy chain. In preliminary experiments *in vitro*, we showed that deletion of the CH1 domain had a negative impact on plasmablast differentiation without affecting other stimulated B cells thus confirming the specific toxicity of truncated/abnormal Ig for PCs as already shown with truncated LCs^30^. Overall, the IgJ^CreERT2^ model appears as an exquisite new tool to study PC development, maintenance and deregulation.

## METHODS

### Generation of transgenic mice

The IgJ-GFP construct includes the 1.1kb NdeI/NheI genomic fragment harboring IgJ enhancer activity previously described^27^ and the 1.4 kb IgJ promoter followed by *EGFP* gene from pMOD-ZGFPsh vector (Cayla Invivogen), all flanked by two insulators from chicken β-globin gene locus [23]. A neomycine resistance cassette follows this construct. CK35 embryonic stem cells were electroporated with the linearized vector and G418 resistant clones were screened by PCR (**supplemental Fig.1**). Two clones (10 and 13) were reimplanted in C57BL/6 blastocysts. For each clone, chimeric founders gave rise to a IgJ^GFP^ lineage respectively named IgJ^GFP^10 and IgJ^GFP^13.

The Jchain^tm1(CreERT2_EGFP)Wtsi^ mice, hereafter called IgJ^CreERT2^, were generated by the EUCOMMTOOLS consortium at the Welcome Trust Sanger Institute of London (MGI:5633773). They were obtained from the French National Infrastructure for Mouse Phenogenomics (Phenomin) at the Mouse clinical institute (Illkirch, France). Briefly, the expression cassette consisting in an eGFP-F2A-Cre^ERT2^ was inserted in the intron between exons 1 and 2 of the J-chain allele. This cassette was preceded by a splice acceptor sequence followed by a F2A sequence allowing the independent expression of *eGFP* and *Cre^ERT2^* genes under the control of the J chain promoter/enhancers. A puromycin resistance cassette flanked by Rox was also inserted in 3’ of the construct. The full construction is depicted in **supplemental Fig.1**.

Rosa26-LSL-Tomato mice were obtained from The Jackson Laboratory (JAX, Bar Harbor, ME, stock number 7914). In a few words, mice contain in the Gt(ROSA)26Sor locus a floxed STOP cassette preventing transcription of a CAG promoter-driven red fluorescent protein variant (tdTomato).

Finally, Heavy Chain Depostion Disease (HCDD)-CH1^+^ mice were generated as previously described^29^. Briefly, the gene coding for a human monoclonal heavy chain obtained from a HCDD patient was introduced into the mouse joining segments in the kappa locus with a floxed CH1 domain.

All the animals were bred and maintained in pathogen-free condition in our animal facility. Unless stated otherwise, heterozygous animals for every transgenic genes were used. All experimental procedures have received the approval of our institutional review board for animal experimentation and of the French Ministry of Research (N° 7655-2016112211028184).

### Immunization and Tamoxifen induction

When needed, mice were immunized with one intra-peritoneal injection of 200μl of sheep red blood cells (Sigma-Aldrich) and analyzed at the indicated times. *In vivo* tamoxifen induction protocol consisted in two intraperitoneal injections of 4mg of Tamoxifen (Sigma-Aldrich) in sunflower oil (Sigma-Aldrich) with a two-day rest. Mice were sacrificed the day after the last tamoxifen injection (**supplemental Fig.2**).

### Flow cytometry and cell sorting

Antibodies and reagents used for staining and cell sorting experiments are detailed in supplemental **Table I**. Flow cytometry analyses were done on a BD Pharmingen LSRFortessa® cytometer. Cell sorting experiments were done on a BD Pharmingen FACSVantage® cell analyzer. Data were then analyzed with BD FACSDiva software (BD Biosciences) and FlowLogic (Miltenyi Biotec).

### Confocal microscopy

For confocal microscopy, organs were incubated 1 h in PBS/PFA 4 % followed by 12h in PBS/sucrose 30 % before freezing. Frozen 8 μm sections were prepared and incubated with PBS/BSA 3 %/Tween 0.05 % before staining. Images were acquired with a Zeiss LSM 510 Meta confocal immunofluorescent microscope (Zeiss) and then analyzed with LSM Image Browser software (Zeiss).

### Spleen cell culture

Total spleen cells were cultured at a density of 1 × 10^6^ cells/mL in RPMI 1640 supplemented with 10% fetal calf serum (FCS) with 5 ng/mL LPS (Invivogen). For 4-hydroxytamoxifen (OHTAM) (Sigma-Aldrich) induction, 1uM were dissolved in methanol were added in the culture media on the third day of culture.

### Screening of CH1 deletion in HCDD-CH1+ mice

Screening of CH1 deletion was carried out using the following primers: pVHForEagI ACGGCCGAAGCTTAAAAACCTCAGAGGATTTGTCATCTCTA and GammaCH3HumRev GTGGTCTTGTAGTTGTTCTC.

### ELISPOT assays

For evaluation of Igκ or IgM secretion, sorted populations were seeded in duplicate at a density starting at 2 × 104 cells per well, followed by five-fold serial dilutions in culture medium on a 96-well plate MultiSreen HTS (Millipore) previously coated over-night at 4 °C with 1.5 μg per well of anti-Igκ or anti-IgM (Beckman Coulter) in PBS and saturated with culture medium. Cells were incubated 6 h at 37 °C and then removed by washing with PBS/Tween 0.01 %. Plate was then incubated 1 h à 37 °C with 1 μg per well of alkaline phosphatase-coupled anti-Igκ or anti-IgM (Beckman Coulter) in PBS. After washing PBS/Tween 0.01 %, plates were incubated 10 min with 100 μL of BCIP/NBT alkalin phosphatase substrate (Millipore). After washing and drying, pictures of wells were taken and images were analyzed for spots numbers and areas with ImageJ software.

### Statistical analysis

The statistical tests used to evaluate differences between variables were indicated in legends and were done using Prism GraphPad software (San Diego, CA). All P values < 0.05 were considered significant using Student’s t-test.

## RESULTS

### Generation and characterization of the IgJ^GFP^ reporter model

To follow *IgJ* expression and create a new reporter model for PC lineage, we generated a transgenic mouse strain that express the eGFP cDNA under the control of regulatory sequences from the IgJ locus (**supplemental Fig.1**). The transgene comprise the 1.4 kb promoter sequence of the *IgJ* gene including the Pax5 repression site ^31^ and a 1.5 kb genomic fragment from a region located between 6 and 7.5 kb upstream of the transcriptional start site, containing the main enhancer element controlling *IgJ* gene expression ^27^. To limit the position effects, the transgene was flanked with two insulators sequences from the chicken β-globin. We obtained two different strains (clone 10 and 13) that were first analyzed by flow cytometry on 8 days SRBC immunized mice. In spleen, we detected two distinct cell populations expressing GFP: a bright population (GFP^high^) characterized by a low expression of the pan-B cell marker B220 and a population expressing intermediate level of GFP (GFP^low^) together with the B220 marker (**Fig.1A**). As expected, in both strains, CD138^+^ B220^low^ PCs were highly GFP positive with a slight difference in intensity between clones 10 and 13 (**Fig.1B**). As clone 13 presented the strongest level of GFP, further results presented in this study will focused on this strain. But, except when indicated, the pattern of expression of the GFP is similar in both strains. We also detected a GFP^high^ population in bone marrow (BM) corresponding to the CD138^+^/B220^low^ PCs and no other GFP^+^ population showing the restriction of *IgJ* expression to the PC compartment. Interestingly, by contrast to previous assumptions ^32^, almost all CD138^+^ cells in spleen or BM were GFP^high^ and thus, express the *IgJ* gene (**Fig.1B**).

**Figure 1:**
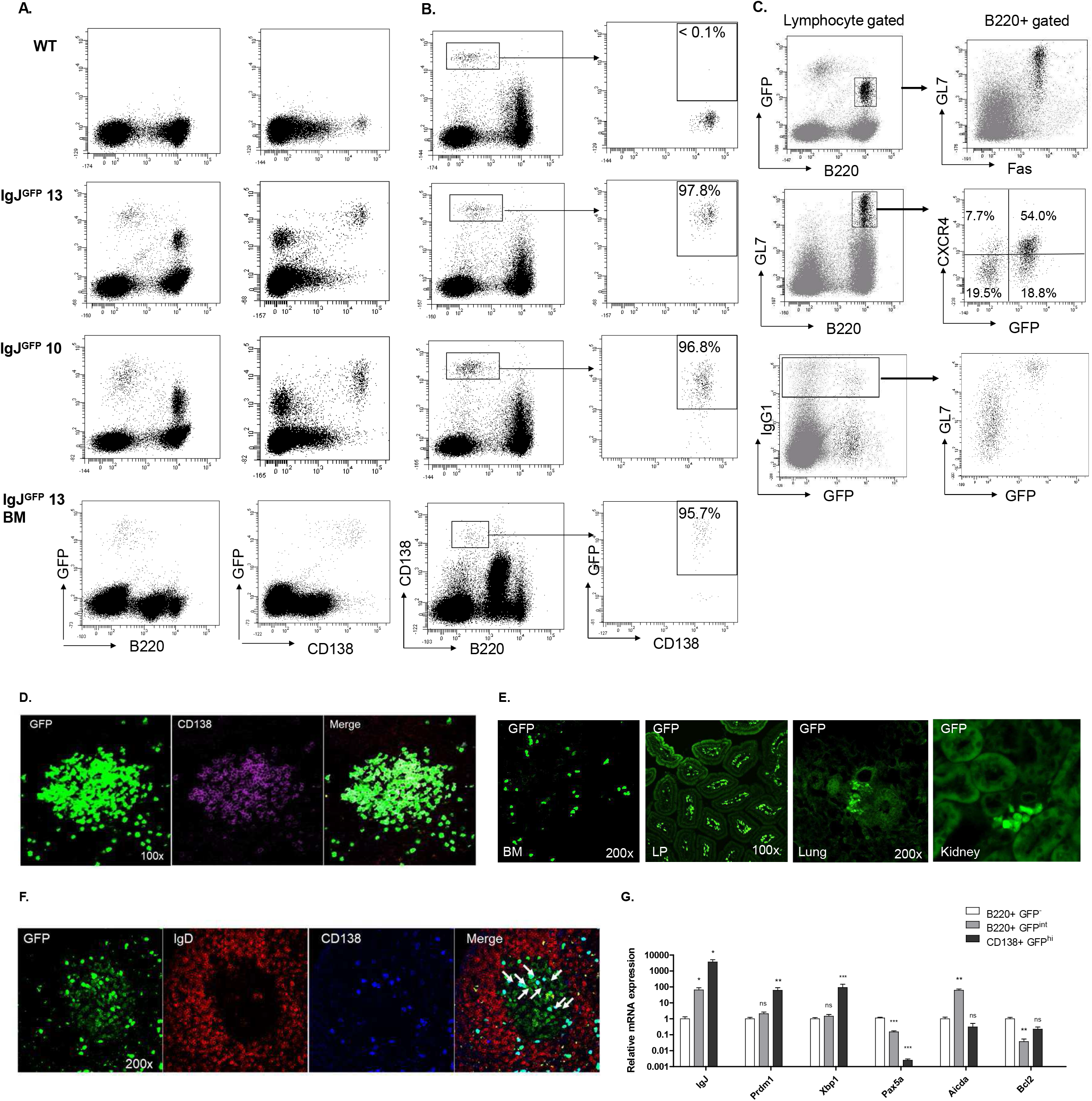
The vast majority of PCs expresses IgJ chain. (A) Flow cytometry analysis of GFP expression in spleen and bone marrow B cells (left) and PCs (right) of IgJ^GFP^ mice. Two distinct populations of GFP expressing cells are observed (B) Flow cytometry analysis of GFP expression among PCs IgJ^GFP^ mice. PCs are gated based on the CD138^+^/B220^low^ staining (left) and analyzed for GFP expression (right). Percentages indicate the proportion of GFP-expressing PCs. Data shown in (A) and (B) are representative of 10 independent experiments with at least 2 mice of each strain. (C) Flow cytometry characterization of the GFP^int^ population in spleen cells of 8 days immunized IgJ^GFP^13 mice. GFP^int^/B220^+^ (upper left panel) cells are framed in the left graph appears in black in the subsequent analysis of GL7 and Fas expression (upper right panel). Analysis of GFP expression among B220^+^/GL7^+^ GC B cells (middle left panel). B220^+^/GFP^int^ cells are framed and appear in black in the subsequent analysis for CXCR4 expression (middle right panel). GL7 and GFP expression in IgG1^+^ B cells is also shown (bottom panels). Percentages of each population are indicated. Data shown are representative of 4 independent experiments with at least 2 mice. (D) Confocal microscopy analysis of spleen section. PC foci in extrafollicular zone from 8 days immunized IgJ^GFP^ mice stained with anti-CD138. Note that almost all PCs coexpress GFP (green) and CD138 (red). (E) GFP^high^ PC are observed in various organs. BM, Bone Marrow; LP, Lamina Propria. (F) GC in a mesenteric lymph node from a SRBC immunized mouse. GFP^low^ cells are in the IgD^neg^ GC area. Note the presence of a significant number of GFP^high^/CD138^+^ cells located within the GC area (arrows). (G) Transcription analysis of sorted B220^+^/GFP^neg^, B220^+^/GFP^int^ and CD138^+^/GFP^high^ cells from spleens of IgJ^GFP^ 13 mice. Fold change expression was established compared to B220^+^/GFP^neg^ B cells with Gapdh as housekeeping gene. Data are pooled from 7 experiments in which spleens from 2 mice were pooled and sorted, except for CD138^+^/GFP^high^ cells that were sorted in 5 experiments. Results are expressed in log scale as mean + SEM.

We also observed a B220^+^/GFP^low^ population that expresses the germinal centers markers GL7 and Fas (Fig. 1C, top panel). About 75% of the GL7^+^ B cells in spleen are GFP positive. Similar results were obtained with PNA/Fas staining in Peyers’ patches (PP) (**supplemental Fig.3**). GFP^low^ GC B cells displayed both centroblast and centrocyte phenotypes as seen with CXCR4 staining (**Fig. 1B**, **middle panel**). Some of the GFP^low^ B cells are IgH-switched expressing IgG1 together with GL7 (**Fig.1C**, **lower panel**). Interestingly, IgH-switched cells that do not express GFP are also GL7^neg^ showing that post-GC memory B cells do not express IgJ (**Fig. 1C**, **lower panel**).

Immunofluorescence on spleen sections confirmed that PCs are almost all GFP^high^ cells, localized mainly in the extrafollicular and peri-arteriolar areas (**Fig.1D**). GFP^high^ cells are readily observed in various organs including BM and lamina propria but also lung or kidney (**Fig.1E**), validating IgJ^GFP^ model as a valuable tool to study PC localization and migration. Consistent with flow cytometry results, GFP^low^ cells colocalized with GC area (IgD^*^) in lymph nodes (**Fig.1F**), spleen and PP and are all GL7^+^ cells (*not shown*). Strikingly, we also observed a significant number of GFP^high^ plasma cells in GCs of spleen and mesenteric lymph nodes of immunized mice (**Fig.1F**). As previously described, some PCs were localized at the periphery of the GCs but we also detected in significant number of GFP^high^ cells within the GC structure. Most of this intra-GC PCs were fully differentiated as seen by their CD138 staining and were mainly located in the light zone (Fig. 1F, right image). To further characterize these IgJ-expressing cells, we performed transcriptional analyses on sorted B220^+^/GFP^low^ and CD138^+^/GFP^high^ cells from spleen relative to B220^+^/GFP^*^ cells. Consistent with their GC phenotype, GFP^low^ B cells expressed high amount of *Aicda* gene together with a significant decrease of the anti-apoptotic transcription factor *Bcl2* compared with GFP^neg^ B cells. As expected, GFP^low^ B cells also expressed increased amount of IgJ transcripts but, interestingly, did not express increased amount of PC transcription factors like *Prdm1* or *Xbp1*. However, they displayed a significant reduction of about 4 fold of *Pax5* (Fig. 1G). Consistent with their PC phenotype, GFP^high^ cells presented with high expression of *IgJ*, *Prdm1*, *Xbp1* together with a strong decrease of *Pax5* (**Fig.1G**).

### IgJ is an early marker of B cell terminal differentiation

To verify if the *IgJ* gene is expressed during all stages of PC differentiation, we further performed *in vitro* LPS stimulations of spleen B cells from IgJ^GFP^ mice. As seen in **Fig.2A**, GFP was strongly detected from the first 48h in a significant proportion of cells, most of these cells remaining CD138^neg^. At 72h, all the CD138^+^ cells were also GFP^+^, but a population of GFP^+^/CD138^neg^ cells was still present. Interestingly, κ light chain specific ELISPOT revealed that those GFP^+^/CD138^neg^ cells are secreting cells even if spots area were slightly smaller than GFP^+^/CD138^+^ plasmablasts cells (**Fig.2B**). By contrast, GFP^neg^/CD138^neg^ B cells secreted no or very low amount of Ig light chain. Real-time PCR analyses confirmed these results showing that GFP^+^/CD138^neg^ cells already express the master regulator of plasma cell *Prdm1* and the secreted form of Igμ heavy chain, together with a slight downregulation of *Pax5* (**Fig.2C**). These results seem to demonstrate that *IgJ* expression appears very early during the process of terminal differentiation of the B cell lineage, before the expression of CD138 and should therefore be an efficient marker of the previously described pre-plasmablast population ^28,33^.

**Figure 2:**
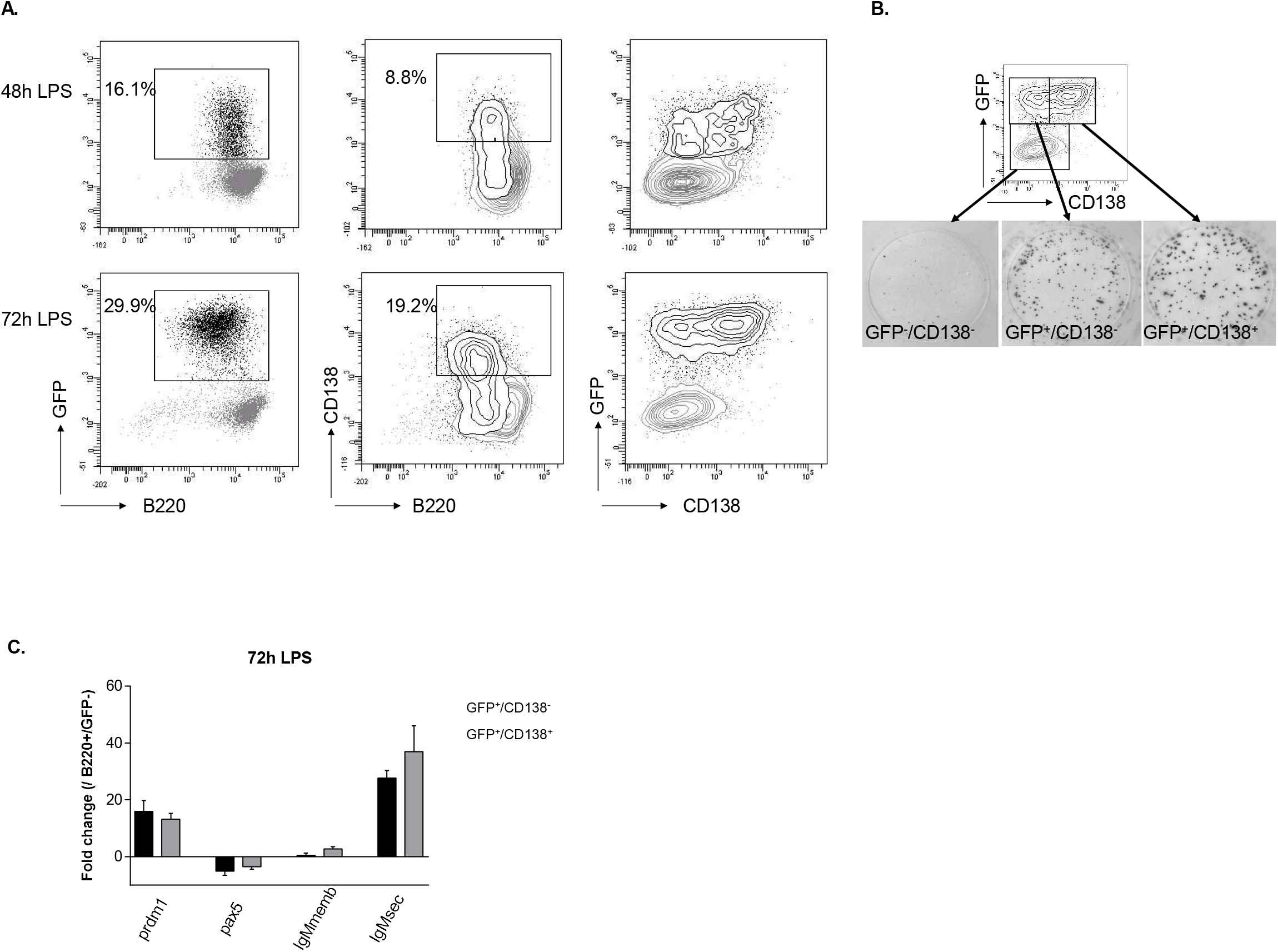
IgJ expression is an early marker of PC differentiation. (A) Flow cytometry analysis LPS-stimulated splenocytes from IgJ^GFP^ 13 mice at 48h and 72h. B220^+^/GFP^int^ cells are framed and percentages are indicated. GFP^+^ population framed in the left panel appears in black in the subsequent graphs. Percentages of CD138^+^ plasmablasts are indicated (middle) and the expression of CD138 among GFP^+^ cells is shown in the right panel. Data shown are representative of 10 independent experiments with at least 2 mice. (B) Representative ELISPOT results from LPS-stimulated cells sorted at 72h according to the GFP and CD138 expression as indicated in images. Data shown are representative of 3 independent sorting experiments. (C) Gene expression relative to *Gapdh* of sorted CD138^neg^/GFP^neg^, CD138^neg^/GFP^+^ and CD138^+^/GFP^+^ cells. Results are shown as fold-change expression compared to the CD138^neg^/GFP^+^ cells. Results are from 3 independent sorting experiments. Data are shown as mean ± SD.

### Recombinase activity in IgJ^CreERT2^ model only targets CD138^+^ cells

Having shown that IgJ promoter/enhancers could be advantageously used to target PC lineage, we characterized the IgJ^CreERT2^ mice line, created by the EUCOMMTOOLS consortium, which carries an eGFP and Cre^ERT2^ recombinase expression cassette under the control of regulatory sequences of the IgJ locus (**supplemental Fig.1**). In contrast with our IgJ^GFP^ model, spleen and BM analysis of these mice revealed a very weak fluorescence of the eGFP making it disadvantageous to follow up PCs and the expression of the Cre^ERT2^ recombinase by flow cytometry (**Fig.3A**). This was confirmed in LPS-stimulated cells that did not show detectable GFP in CD138^+^ plasmablasts (**Fig.3B**).

**Figure 3:**
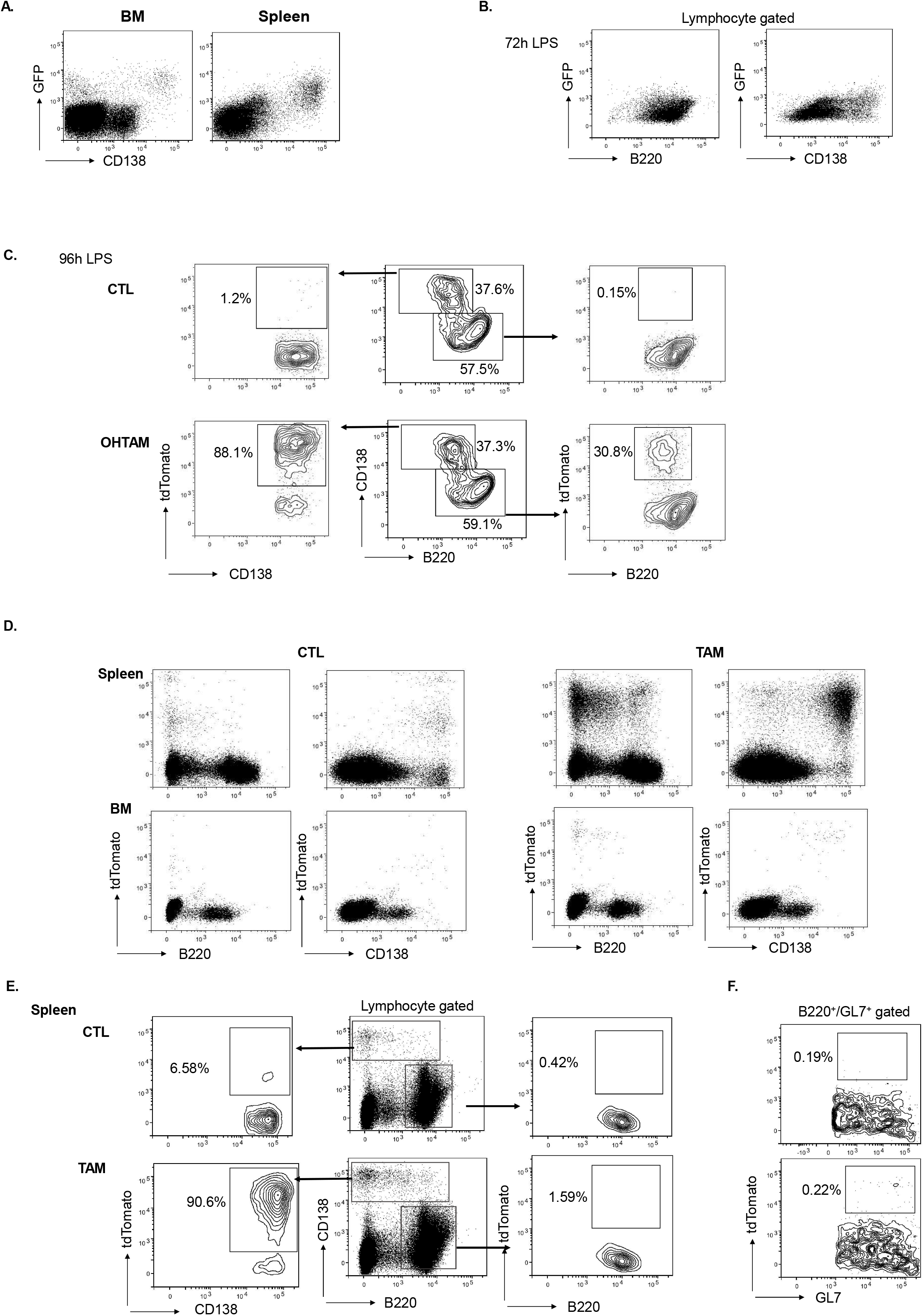
Recombinase activity in IgJ^CreERT2^ targets CD138^+^ cells. (A) Flow cytometry analysis of eGFP expression in spleen and bone marrow PCs of IgJ^CreERT2^ mice denoting the low fluorescence (B) Flow cytometry analysis LPS-stimulated splenocytes at 72h for eGFP and B220^+^ or CD138^+^ (C) Flow cytometry analysis LPS-stimulated splenocytes from IgJ^CreERT2^ x tdTomato mice at 96h. tdTomato expression in B cells (right) and PCs (left) for the control (CTL) vs 4-hydroxytamoxifen (OHTAM) treated cells. Cells expressing tdTomato are mostly found in the OHTAM group. As previously shown in IgJ^GFP^ mice, some CD138^−^ cells already expressed tdTomato (right). (D) Representative flow cytometry analysis for tdTomato expression in spleen and bone marrow of IgJ^CreERT2^ x tdTomato mice after *in vivo* treatment. (E) Representative flow cytrometry analysis of spleen cells for tdTomato expression. PCs are gated based on the CD138^+^/B220^neg/low^ staining (middle) and analyzed for tdTomato expression (left). B cells are gated based on B220^high^/CD138^neg/low^ (middle) and analysed for tdTomato expression (right). Percentages indicate the proportion of tdTomato-expressing cells. (F) Flow cytometry analysis of B220^+^/GL7^+^ GC B cells showing almost no expression of tdTomato.

To further characterize the IgJ^CreERT2^ recombinase activity, mice were crossed with the Rosa26-LSL-Tomato mouse line (tdTomato) in which a loxP-flanked STOP cassette prevents transcription of the downstream red fluorescent protein tdTomato. Total spleen cells from IgJ^CreERT2^ x tdTomato mice, were put in culture under LPS stimulation and treated with OHTAM after 3 days of culture.

On the following day (96h of culture), we achieved 77.13% ± 8.23 (mean ± SEM n=3) of CD138^high^/tdTomato^+^ cells while the B220^high^/CD138^neg^ remained low (8.51% ± 2.95 (mean ± SEM n=3)). However, about 25% of B220^low^/CD138^neg^ cells acquired the tdTomato fluorescence. These cells likely correspond to the pre-plasmablasts described in our IgJ^GFP^ model, that expressed *IgJ* gene before the expression of CD138. There is no leaky Cre activity in the stimulated cells not treated with OHTAM (**Fig3.C**).

*In vivo*, the IgJ^CreERT2^ x tdTomato offspring was immunized with SRBC and treated with tamoxifen (**supplemental Fig.2**). At day 11 post-immunization, following 2 injections of tamoxifen, we detected the presence of tdTomato in 91.04% ± 1.14 (mean ± SEM n=3) of B220^low^/CD138^high^ cells in spleen (**Fig.3D**) and 86.19% ± 6.49 (mean ± SEM) in BM. Less than 1% of B220^high^/CD138^neg^ B-cells in the spleen were tdTomato^+^ likely accounting for few B cells already committed to the PC fate. No other cells were tdTomato^+^ demonstrating a high specificity of the IgJ activity as seen in IgJ^GFP^ mice (**Fig.3E**). In contrast with the IgJ^GFP^ model, we did not detected Cre^ERT2^ activity in GC B cells (**Fig.3F**). This discrepancy is likely due to the weak expression of Cre^ERT2^ in these cells as shown by the absence of the coexpressed GFP. In control mice injected with oil alone, we detected a leaky activity of the Cre^ERT2^ recombinase in PCs (10.25% ± 5.51 mean ± SEM; n=3 in the spleen and 22.56% ± 12.30 mean ± SEM; n=3 in BM), that was gender independent. Such leaky activity of the Cre^ERT2^ recombinase has already been described in other models^34,35^ and will need to be taken in account for genetic manipulations of PCs in this model. Altogether, our data show that genetic manipulations of PCs can be efficiently and specifically obtained in IgJ^CreERT2^ mice.

### Modifications affecting immunoglobulin structure affect PC survival

To further validate the IgJ^CreERT2^ model as an efficient tool to study PCs biology, we took advantage of our previously published mouse model for Heavy Chain Deposition Disease (HCDD)^29^ in which the gene coding for a pathogenic human Igγ heavy chain was inserted in the Igκ locus of the mice. The CH1 domain of this HC was flanked by two loxP sites to allow its deletion upon Cre recombinase activity since the disease is characterized by deposition of a truncated CH1-deleted HC. CH1 domain deletion was not supposed to affect PCs since CH1-deleted HCs do not interact with the binding immunoglobulin protein (BiP) chaperone and were then supposed to be freely secreted ^36,37^. In our previous study, the deletion of the CH1 was done in the germline lineage so that all newly formed B cells expressed the truncated HC. While we did not detected any effect on the PC differentiation, we showed that the truncated HC was poorly secreted compared to the full-length and that PCs bearing this truncated Ig were more sensitive to proteasome inhibitors and displayed increased ER stress markers^29^. Therefore, to assess the real effect of deleting the CH1 only in PCs, we crossed the IgJ^CreERT2^ mouse line with the HCDD-CH1^+^ mouse, carrying a floxed CH1 domain. In a first validating experiment, we sorted spleen cells to obtain the following populations: plasma cells (B220^low^/CD138^+^ cells), germinal center B cells (B220^+^/GL7^+^), other B cells (B220^+^) and the rest of cells. Cells were cultured without stimulation for 24h with OHTAM before gDNA from each population was extracted to carry out a PCR allowing the detection of the deleted CH1 allele. As seen in **Fig.4A**, the CH1-deleted band appeared only in the PC population. In contrast with the tdTomato *in vivo* experiment, we did not observe any leaky activity *in vitro* for this model as the non-treated PCs only displayed the band corresponding to the full-length HC. Then, total spleen cells from IgJ^CreERT2^ x HCDD-CH1^+^ mouse were stimulated with LPS for 4 days and treated with OHTAM or vehicle from day 3. We first confirmed that OHTAM treatment induced the deletion of the CH1 exon with no leaky activity in the non-treated cells (**Fig.4B**). As seen in **Fig.4C**, we observed a decreased of CD138^+^/B220^low^ cells treated with OHTAM compared to non-treated ones (32.91% ± 2.91 vs. 23.82% ± 4.54 mean ± SEM, \**P* <0.05; n=3) showing that truncation of the CH1 domain could be toxic when occurring in PCs.

**Figure 4:**
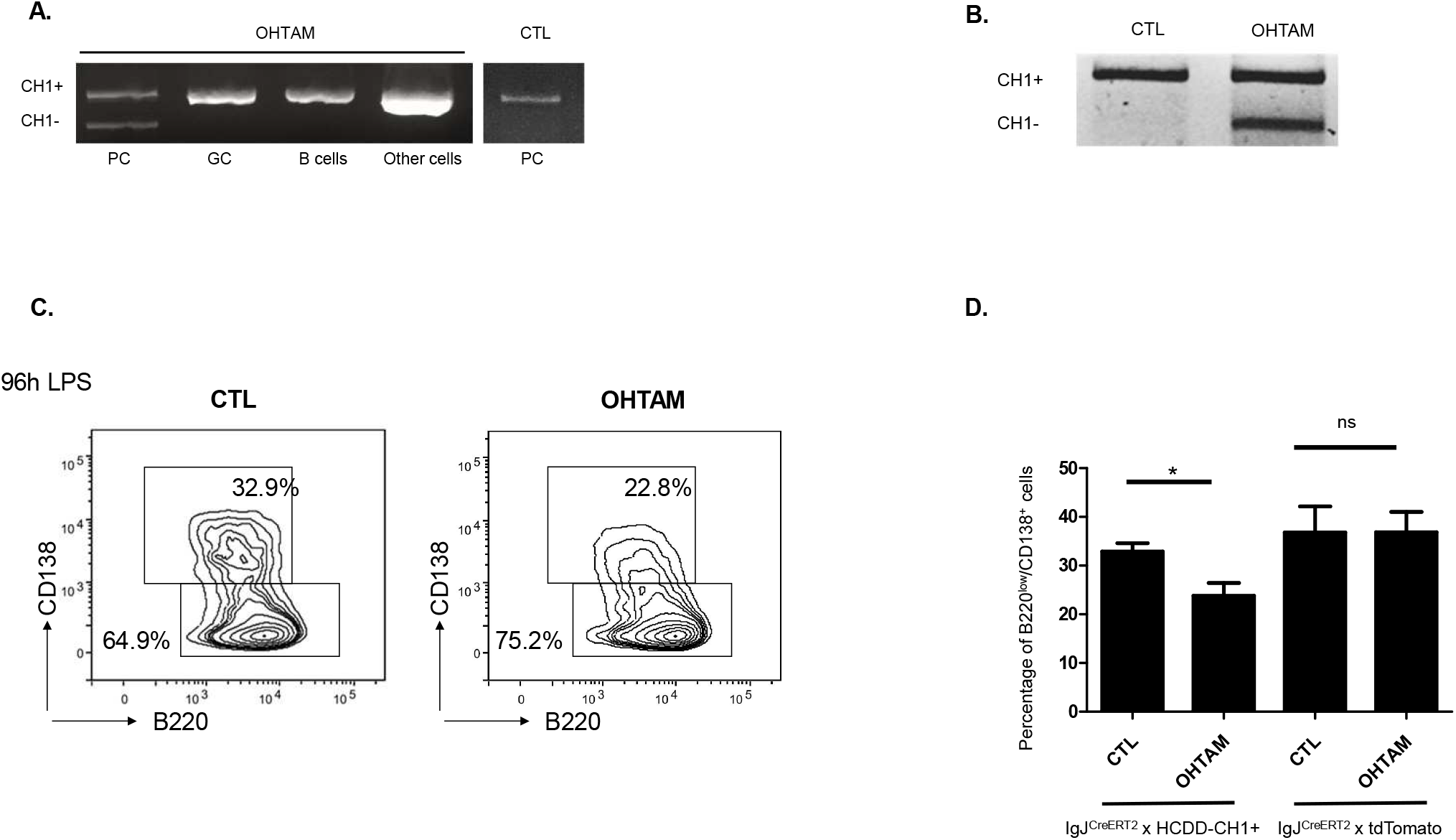
PC survival correlates with Immunoglobulin modification. (A) genomicDNA analysis by PCR of sorted spleen cells of IgJ^CreERT2^ x HCDD-CH1^+^ mouse (Plasma cells : PC ; Germinal Center cells: GC, B cells and rest of cells) after being cultured for 24h with OHTAM. CTL non-TAM treated PCs do not exhibit the truncated band (right). (B) genomicDNA analysis by PCR of 4 days LPS-stimulated splenocytes from IgJ^CreERT2^ x HCDD-CH1^+^ mice to show the CH1 deletion only in OHTAM cells cultured with it for 24h. (C) Representaive flow cytometry analysis of 4 days LPS-stimulated splenocytes from IgJ^CreERT2^ x HCDD-CH1^+^ after 24h of treatment with gates showing the percentage of CD138^+^ cells and B220^+^ cells. (D) Histograms showing the significant decrease of plasmablasts (B220^low^/CD138^+^ population) following the deletion of CH1 in IgJ^CreERT2^ x HCDD-CH1^+^ mice splenocytes. IgJ^CreERT2^ x tdTomato mice were used as control, showing that this decrease is not due to OHTAM treatement. ns, non significant; *P < 0.05.

## DISCUSSION

This study sheds light on a yet unknown pattern of expression of the IgJ chain in mouse. The lack of reliable antibodies has long precluded the efficient characterization of the IgJ-expressing cells in mouse. Here, using a transgenic reporter system in which the GFP gene was placed under the control of the IgJ promoter/enhancer, we show that IgJ chain is expressed in almost all PCs in mouse, independently of their localization in spleen, BM, lamina propria or PP. Our transgenic model allowed us to precisely determine the stages of induction of the IgJ chain *in vitro*, following LPS stimulation of primary B cells, with IgJ/GFP expressed earlier than CD138. This result could be consistent with preplasmablasts (PPBs), an early subset of ASCs described by Kallies et al. ^28^. Using mice rendered prdm1-deficient by the targeted insertion of a GFP gene, they characterized a population of low secreting cells that express IgJ together with markers of PC differentiation like XBP1 and IRF4. However, the GFP^+^/CD138^neg^ cells in our study secrete high amounts of Ig, comparable to CD138^+^ cells. This difference could account for the absence of BLIMP1 in their model, limiting the secretion rate of Ig chains ^22^. Whether PPBs in their study and our GFP^+^/CD138^neg^ cells represent different stages of PC differentiation or an equivalent population, remains to be addressed. In any case, IgJ expression appears as a reliable marker of ASCs *in vitro* as already shown in human B cells ^23,38,39^ and the IgJ^GFP^ model will be useful to accurately study the early steps of the PC differentiation. *In vivo*, we confirmed than no other hematologic cell lineage than the B lineage expressed the *IgJ* gene, making it a better reporter than other PC-related genes like Blimp-1^19,18^, which are expressed in other cell lineages. Interestingly, we did not detected any IgJ activity in the pre-B and immature B cell stages as previously claimed ^24^. However, we did detect IgJ activity in GC B cells. This activity could mirror cells that are already engaged in the PC differentiation. However, our observations showed that most GC B cells were GFP^+^ and all of them cannot be committed to PC differentiation. In fact, GFP expression in these cells could reflect a transient activity of the *IgJ* gene in GCs that, together with the long half-life of the GFP protein^40^, could be misleading on the real activity of the IgJ promoters. More experiments are needed to conclude on this expression of the *IgJ* gene in GC and could elucidate new early steps of PC differentiation.

Since the IgJ^GFP^ model put light in the use of IgJ promoter as a good one to target PCs, the creation of a mouse model having an inducible recombinase under its control seemed like the path to take to improve, not only the study of the effects of modifying PC but the creation of mouse models for several PC-related malignancies. Here we presented the *in vitro/in vivo* characterization of the inducible IgJ^CreERT2^ inducible model established by the EUCOMM consortium. As discussed above, this model also expressed an *eGFP* gene together with the tamoxifen-inducible Cre^ERT2^ recombinase. Contrastingly to IgJ^GFP^, eGFP levels were very weak. We attributed this discrepancy to the complex architecture of the IgJ^CreERT2^ locus (artificial splice donor site, two consecutive T2A cleavage) that likely affects the global expression of the transgenes. However, we cannot exclude that the transgenic construct of the IgJ^GFP^ mice, which is not a knock-in and does not contain the full DNA region surrounding *IgJ* gene, could also be a bit further from the physiologic regulation of the locus. The low level of fluorescence of the eGFP in the IgJ^CreERT2^ model could be improved by the use of anti-GFP antibodies in flow cytrometry to correlate the expression of Cre^ERT2^ recombinase in future models if other reporter genes are not available. Nonetheless, with the reporter tdTomato model, we showed that the recombinase activity only targets CD138^+^ cells while B cells remain almost untouched. However, the leakiness of Cre recombinase in PCs (non subjected to tamoxifen) could be more worrying for temporal induction of oncogenes for example. This leaky affect has been well described in the literature ^34 35^. Reaching no-leak seems almost impossible, but it could likely be improved by separating the treated mice from those that are not to avoid the cross-contamination by coprophagia^34^. In any case, if temporal activity of the Cre recombinase is not fully ensured in this model, its spatial one seems to be high specific with a strict restriction of activity to antibody secreting cells and for the first time, no other hematopoietic cell lineage.

Our preliminary results with the HCDD-CH1^+^ model showed that PC survival is linked to the maintenance of its Ig integrity. This result demonstrated that even if CH1 domain is not strictly required for HC to be secreted^36^, its deletion seems to affect PC survival. The absence of truncated band in untreated cells that would appear due to the leaky activity of the recombinase could be explained by this deleterious effect: those plasmablasts that spontaneously delete their CH1 domain are automatically eliminated during differentiation into ASCs and therefore cannot be retrieved by PCR. These results could account for the rare occurrence of heavy chain diseases (HCD) or heavy chain deposition diseases (HCDD) characterized by the production of CH1-truncated HCs^41,37^. Moreover, the rare HCDD patients are mostly characterized by small, low-proliferative PC clones and low levels of circulating monoclonal HC. We have also shown in our mouse model of HCDD that the isolated truncated HC is far less secreted than the full-length HC associated with LC, highlighting that the CH1 domain, if not mandatory, at least greatly improved Ig trafficking into PCs^29^. This low secretion was also associated with increased ER stress and higher sensitivity to proteasome inhibitors^29^. Altogether, these previous findings and observations represented a body of evidence that CH1 deletion was somehow embarrassing for secreting cells. However, since CH1 deletion in these cases appears early in B cell development (mouse model) or is associated with oncogenic events (HCDD patients), it remained difficult to evaluate the real toxicity of a truncated HC for PCs. We now showed that CH1 deletion directly occurring in PCs is highly toxic, as previously shown in our laboratory for truncated LCs^30^, paving the way for new therapeutics targeting monoclonal Ig. Currently, studies using of Antisense Oligo Nucleotides (ASOs) to induce exon skipping of Ig genes are being developed and could be adapted to be used in PC malignancies to generate toxic truncated Igs ^42^.

Altogether, we herein described two new models to study PCs using the restricted expression of *IgJ* gene to antibody secreting cells. IgJ^GFP^ mice will reinforce the pool of reporter models for PCs with a more specific pattern of expression. The use of IgJ^CreERT2^ model will be a huge step in the understanding of PC physiology and the modelling of PC neoplasm. The breeding of this novel inducible model with others carrying Cre-inducible genes or oncogenes opens new avenues to study this crucial but yet unexplored lineage.

## ACKNOWLEDGMENTS

The authors thank the staff of the BISCEm technical platforms at the University of Limoges.

## Notes

**Conflict of Interest:** The authors declare that the research was conducted in the absence of any commercial or financial relationships that could be construed as a potential conflict of interest

**FUNDING** This work was supported by grants from Fondation Française pour la Recherche contre le Myélome et les Gammapathies monoclonales, Limousin committees of Ligue nationale contre le cancer, Agence régionale de la santé and Institut Universitaire de France. MVA was funded by fellowships from Région Limousin (now Région Nouvelle Aquitaine) and Fondation ARC pour la Recherche sur le Cancer. BS is supported by Centre Hospitalier Universitaire Dupuytren Limoges and Plan National Maladies Rares. FL and AB were funded by French government fellowships. JML was funded by French government fellowships and Ligue national contre le cancer.

### Competing Interest Statement

The authors have declared no competing interest.

## REFERENCES

1. Delogu, A. et al. Gene repression by Pax5 in B cells is essential for blood cell homeostasis and is reversed in plasma cells. Immunity 24, 269–281 (2006).

2. Nera, K.-P. et al. Loss of Pax5 promotes plasma cell differentiation. Immunity 24, 283–293 (2006).

3. Tunyaplin, C. et al. Direct repression of prdm1 by Bcl-6 inhibits plasmacytic differentiation. J. Immunol. Baltim. Md 1950 173, 1158–1165 (2004).

4. Muto, A. et al. Bach2 represses plasma cell gene regulatory network in B cells to promote antibody class switch. EMBO J. 29, 4048–4061 (2010).

5. Sciammas, R. et al. Graded expression of interferon regulatory factor-4 coordinates isotype switching with plasma cell differentiation. Immunity 25, 225–236 (2006).

6. Klein, U. et al. Transcription factor IRF4 controls plasma cell differentiation and class-switch recombination. Nat. Immunol. 7, 773–782 (2006).

7. Shaffer, A. L. et al. Blimp-1 orchestrates plasma cell differentiation by extinguishing the mature B cell gene expression program. Immunity 17, 51–62 (2002).

8. Lin, K.-I., Angelin-Duclos, C., Kuo, T. C. & Calame, K. Blimp-1-dependent repression of Pax-5 is required for differentiation of B cells to immunoglobulin M-secreting plasma cells. Mol. Cell. Biol. 22, 4771–4780 (2002).

9. Reimold, A. M. et al. Plasma cell differentiation requires the transcription factor XBP-1. Nature 412, 300–307 (2001).

10. Fiancette, R. et al. A myeloma translocation-like model associating CCND1 with the immunoglobulin heavy-chain locus 3’ enhancers does not promote by itself B-cell malignancies. Leuk. Res. 34, 1043–1051 (2010).

11. Katz, S. G. et al. Mantle cell lymphoma in cyclin D1 transgenic mice with Bim-deficient B cells. Blood 123, 884–893 (2014).

12. Knittel, G. et al. B-cell-specific conditional expression of Myd88p.L252P leads to the development of diffuse large B-cell lymphoma in mice. Blood 127, 2732–2741 (2016).

13. Dogan, I. et al. Multiple layers of B cell memory with different effector functions. Nat. Immunol. 10, 1292–1299 (2009).

14. Wen, Z. et al. Expression of NrasQ61R and MYC transgene in germinal center B cells induces a highly malignant multiple myeloma in mice. Blood (2020) doi:10.1182/blood.2020007156.

15. Shaffer, A. L., Emre, N. C. T., Romesser, P. B. & Staudt, L. M. IRF4: Immunity. Malignancy! Therapy? Clin. Cancer Res. Off. J. Am. Assoc. Cancer Res. 15, 2954–2961 (2009).

16. Linterman, M. A. & Vinuesa, C. G. Signals that influence T follicular helper cell differentiation and function. Semin. Immunopathol. 32, 183–196 (2010).

17. Fairfax, K. A. et al. Different kinetics of blimp-1 induction in B cell subsets revealed by reporter gene. J. Immunol. Baltim. Md 1950 178, 4104–4111 (2007).

18. Fooksman, D. R. et al. Development and migration of plasma cells in the mouse lymph node. Immunity 33, 118–127 (2010).

19. Kallies, A. et al. Plasma cell ontogeny defined by quantitative changes in blimp-1 expression. J. Exp. Med. 200, 967–977 (2004).

20. Linterman, M. A. et al. Foxp3+ follicular regulatory T cells control the germinal center response. Nat. Med. 17, 975–982 (2011).

21. Xin, A. et al. A molecular threshold for effector CD8 + T cell differentiation controlled by transcription factors Blimp-1 and T-bet. Nat. Immunol. 17, 422–432 (2016).

22. Tellier, J. et al. Blimp-1 controls plasma cell function through the regulation of immunoglobulin secretion and the unfolded protein response. Nat. Immunol. 17, 323–330 (2016).

23. Koshland, M. E. The coming of age of the immunoglobulin J chain. Annu. Rev. Immunol. 3, 425–453 (1985).

24. Max, E. E. & Korsmeyer, S. J. Human J chain gene. Structure and expression in B lymphoid cells. J. Exp. Med. 161, 832–849 (1985).

25. Bjerke, K. & Brandtzaeg, P. Terminally differentiated human intestinal B cells. J chain expression of IgA and IgG subclass-producing immunocytes in the distal ileum compared with mesenteric and peripheral lymph nodes. Clin. Exp. Immunol. 82, 411–415 (1990).

26. Brandtzaeg, P. & Johansen, F.-E. Mucosal B cells: phenotypic characteristics, transcriptional regulation, and homing properties. Immunol. Rev. 206, 32–63 (2005).

27. Kang, C. J., Sheridan, C. & Koshland, M. E. A stage-specific enhancer of immunoglobulin J chain gene is induced by interleukin-2 in a presecretor B cell stage. Immunity 8, 285–295 (1998).

28. Kallies, A. et al. Initiation of plasma-cell differentiation is independent of the transcription factor Blimp-1. Immunity 26, 555–566 (2007).

29. Bonaud, A. et al. A mouse model recapitulating human monoclonal heavy chain deposition disease evidences the relevance of proteasome inhibitor therapy. Blood 126, 757–765 (2015).

30. Srour, N. et al. A plasma cell differentiation quality control ablates B cell clones with biallelic Ig rearrangements and truncated Ig production. J. Exp. Med. 213, 109–122 (2016).

31. Wallin, J. J. et al. B cell-specific activator protein prevents two activator factors from binding to the immunoglobulin J chain promoter until the antigen-driven stages of B cell development. J. Biol. Chem. 274, 15959–15965 (1999).

32. Erlandsson, L. et al. Joining chain-expressing and -nonexpressing B cell populations in the mouse. J. Exp. Med. 194, 557–570 (2001).

33. Jourdan, M. et al. Characterization of a transitional preplasmablast population in the process of human B cell to plasma cell differentiation. J. Immunol. Baltim. Md 1950 187, 3931–3941 (2011).

34. Kristianto, J., Johnson, M. G., Zastrow, R. K., Radcliff, A. B. & Blank, R. D. Spontaneous recombinase activity of Cre–ERT2 in vivo. Transgenic Res. 26, 411–417 (2017).

35. Álvarez-Aznar, A. et al. Tamoxifen-independent recombination of reporter genes limits lineage tracing and mosaic analysis using CreERT2 lines. Transgenic Res. 29, 53–68 (2020).

36. Feige, M. J., Hendershot, L. M. & Buchner, J. How antibodies fold. Trends Biochem. Sci. 35, 189–198 (2010).

37. Bridoux, F. et al. Unravelling the immunopathological mechanisms of heavy chain deposition disease with implications for clinical management. Kidney Int. 91, 423–434 (2017).

38. Brandtzaeg, P. Immunohistochemical characterization of intracellular J-chain and binding site for secretory component (SC) in human immunoglobulin (Ig)-producing cells. Mol. Immunol. 20, 941–966 (1983).

39. Hajdu, I., Moldoveanu, Z., Cooper, M. D. & Mestecky, J. Ultrastructural studies of human lymphoid cells. mu and J chain expression as a function of B cell differentiation. J. Exp. Med. 158, 1993–2006 (1983).

40. Corish, P. & Tyler-Smith, C. Attenuation of green fluorescent protein half-life in mammalian cells. Protein Eng. 12, 1035–1040 (1999).

41. Cogné, M., Preud’homme, J. L. & Guglielmi, P. Immunoglobulin gene alterations in human heavy chain diseases. Res. Immunol. 140, 487–502 (1989).

42. Ashi, M. O. et al. Physiological and druggable skipping of immunoglobulin variable exons in plasma cells. Cell. Mol. Immunol. 16, 810–819 (2019).

