## supplemental Fig. for "New models to study plasma cells in mouse based on the restriction of IgJ expression to antibody secreting cells"

**Supp. Figure 1:** (A) Representation of the IgJ$^{GFP}$ mouse transgene for the expression eGFP (not in scale). (B) Representation of the IgJ$^{CreERT}$ mouse insert for eGFP and Cre$^{ERT2}$ recombinase expression in the mouse IgJ locus (not in scale). I, intron; PA, polyadenylation site; En2SA, En2 splice acceptor; F2A, foot-and-mouth disease virus 18 self-cleaving peptide

**Supp. Figure 2:** Protocol applied for tdTomato expression induction in vivo in IgJ$^{CreERT}$ x tdTomato mice.

**Supp. Figure 3: PNA$^{+}$/Fas$^{+}$ GC B cells in Peyers' patches express the IgJ chain**. Flow cytometry analysis of PNA$^{+}$/Fas$^{+}$ B cells according to their expression of GFP in IgJ$^{GFP}$ mice. Percentages indicate the proportion of GFP-expressing B cells among the population of GC B cells in Peyers' patches. Data shown are representative of 5 independent experiments.

### Supp. Figure 1

**A.**

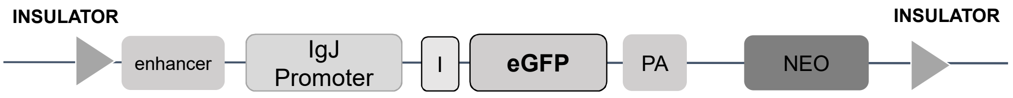


**B.**

#### IgJ locus

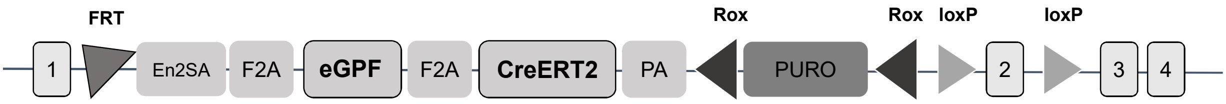


### Supp. Figure 2

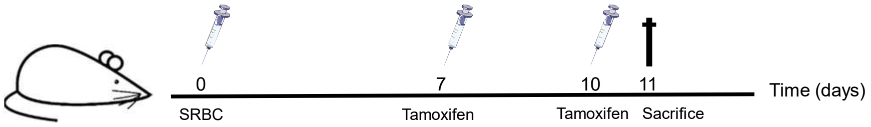


### Supp. Figure 3

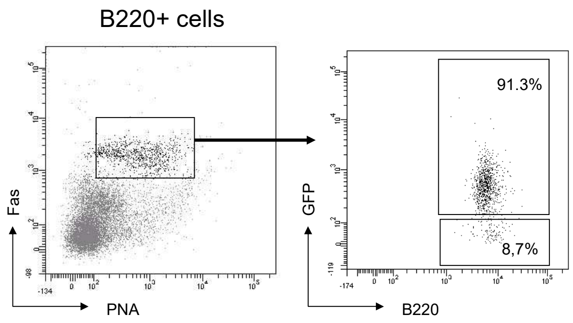

**Supplemental Table. Antibodies used in flow cytometry and cell-sorting experiments**

| Antibody | Clone | Source |
|---|---|---|
| Anti-B220- BD Horizon™ V450 | RA3-6B2 | BD Biosciences |
| Anti-B220-APC | RA3-6B2 | BioLegend |
| Anti-CD138-PE | 281-1 | BD Biosciences |
| Anti-CD138-APC | 281-2 | BD Biosciences |
| Anti-IgD-APC | 11-26C.2a | BD |
| Anti-IgJ-PEa | NA | Santa Cruz Biotechnology |
| GL7-Alexa Fluor®647 | GL7 | BD Pharmingen |
| GL7-PEa | GL7 | BD Pharmingen |
| Anti-CXCR4-PE | 2B11 | BD Pharmingen |
| Anti-IgG1-PE | A85-1 | BD Pharmingen |
| Anti-Fas | JO2 | BD Pharmingen |
| Anti-CD138-BV421 | 281-2 | BD Biosciences |
| Anti-B220-BV786 | RA3-6B2 | BD Biosciences |
